# Molecular characterization and functional annotation of a hypothetical protein (TDB29877.1) from probiotic bacteria *Lactobacillus acidophilus*: an *in-silico* approach

**DOI:** 10.1101/2021.07.23.453527

**Authors:** Md Ataul Goni Rabbani

**Affiliations:** Poultry Production Research Division, Bangladesh Livestock Research Institute (BLRI), Savar, Dhaka, Bangladesh

**Keywords:** *Lactobacillus acidophilus*, stress-response, probiotic, functional annotation, energy minimization, defense mechanism

## Abstract

*Lactobacillus acidophilus* bacteria are widely used as probiotic and to produce various healthy fermented food products. The PNW3 strain of the bacteria has numerous proteins in its genome and some are considered as hypothetical proteins. The aim of the present study was to predict the structures and biological functions of the hypothetical protein (accession number: TDB29877.1) from *L. acidophilus* through an *in-silico* approach applying various bioinformatics tools. The sequence similarity was searched on the available biological databases using BLASTp program to find out the homologues proteins. Besides, determination of various properties like physicochemical characteristics, subcellular localization, phylogenetic analysis, functional annotation, protein-protein interaction, determination of secondary and tertiary structures, active site detection and further quality assessment analysis were done using appropriate computational methods of bioinformatics. *In-silico* analysis revealed that the hypothetical protein has contained TerB-N and TerB-C domains with the presence of YjbR-like superfamily. The Protein-protein interaction network analysis revealed that the protein highly interacted with various known and unknown proteins responsible for iron ion binding, DNA and RNA metabolisms and numerous repair mechanisms that maintain cellular integrity. It was also found that the protein has predominantly alpha-helices in its secondary structure and the three dimensional model has been found to be novel as it possessed expected quality while gone through various quality assessment tools. Thus, the present result indicated that the selected hypothetical protein which is cytoplasmic in nature with Belta-grasp fold, plays important role in responding during stress condition or phage defense mechanism.

## Introduction

*Lactobacillus acidophilus* bacteria are widely used as probiotic for commercial use in the livestock sector especially in poultry feed as additive for the improvement of production performance [1–6]. In addition to produce various healthy food products like yogurt, cheese, and other dairy products [7, 8]. Anyway, probiotics are the direct fed micro-organisms which when administered in adequate amounts confer a health benefit on the host and by improving its intestinal microbial balance [9–11]. These beneficial properties of *L. acidophilus* bacteria has aroused interest of the researchers across the globe to investigate the function of different proteins involved in defensing mechanism.

Advancement in computational biology, development of various bioinformatics tools and analysis servers make it easier to predict functions of the protein, identify sequence similarity, conduct phylogenetic analysis, evaluate active site residue similarity, protein– protein interaction, conserved domains, motif phosphorylation regions and so on [12–17]. In most of the completely sequenced genomes, almost 60% genes have known functions. The number of genes having unknown functions called hypothetical proteins [18] which can be classified as either uncharacterized protein families or domain of unknown functions [19]. Besides, with the advancement in sequencing technologies, the number of sequences deposited in the public biological databases like Swiss-Prot or GenBank have been increasing day by day [20, 21] in comparison to experimentally determined structures deposited in the Protein Data Bank (PDB) [22]. This is resulting a gap between the number of known sequences and confirmed functions. The *in-silico* approaches can minimize the gap predicting structures and biological functions of the proteins [23]. The development of various bioinformatics tools come as a boon in this regard [24] which may also help in designing potential drug against pathogenic organisms and confer efficient pharmacological targets [25, 26].

The annotation report from NCBI-Genome (http://www.ncbi.nlm.nih.gov/genome/) stated that the *Lactobacillus* bacteria have over 50 species. The PNW3 strain of *L. acidophilus* is gram-positive and has a total of 1776 proteins of which 255 proteins are uncharacterized and termed as hypothetical proteins. As the structures and biological functions of these hypothetical proteins are yet to known, molecular characterization and functional annotation of such proteins can lead the relevant researchers to a new dimension of knowledge about the structures, pathways, functions and potential uses in different areas of science. Tremendous development of *in-silico* analysis, numerous bioinformatics tools make it easier to analyze functional annotation of those hypothetical proteins. Thus, the present study was designed to reveal molecular characterization and functional annotation of a hypothetical protein (accession number: TDB29877.1) of the important probiotic bacteria *L. acidophilus* for better understanding of the protein applying various bioinformatics tools.

## Materials and methods

### Retrieval of hypothetical protein sequence

The sequence information of the hypothetical protein from *L. acidophilus* organism (TDB29877.1) was retrieved from the NCBI-Protein database [8]. Then, the sequence was stored as a FASTA format sequence for further use.

### Physicochemical properties analysis

The physicochemical properties of the uncharacterized protein were obtained using the PortParam tool of the ExPaSy server [27]. Various parameters like the molecular weight, amino acid composition, atomic composition, estimated half-life, theoretical pI, extinction coefficient, instability index, aliphatic index, grand average of hydropathicity (GRAVY) etc were analyzed using the tool.

### Homology identification

Similarity search for finding homologous proteins from related organisms that might have structural similarities with the selected hypothetical protein was carried out using BLASTp program of NCBI against non-redundant protein sequences and UniProt databases [28–30].

### Multiple sequence alignment and phylogeny analysis

Multiple sequence alignment (MSA) was done by MUSCLE using MEGA software [31] between the selected hypothetical protein and other proteins obtained from BLASTp program of NCBI. MSA was also cross-checked by Clustal Omega program of EMBL-EBI [32]. Then, Phylogeny.fr tool [33] was used for the phylogeny analysis of the selected protein sequences.

### Subcellular localization analysis

The subcellular localization of the selected hypothetical protein was predicted by using CELLO server [34]. PSORTb [35] and SOSUI tool [36] were also used for the verification of the subcellular localization. In addition, TMHMM, HMMTOP and CCTOP tools [37–39] were also used to cross-check the results.

### Functional annotation analysis

For the purpose of functional annotation analysis of the selected hypothetical protein, search carried out at Conserved Domain Database (CDD) of NCBI for conserved domain(s) [40, 41]. Motif search were carried out using Motif server [42] and ScanProsite tool of ExPasY server [43]. Pfam [44] and Superfamily [45] database searches were also done to assign the protein’s evolutionary relationships. InterProScan [46] was employed for the functional analysis of the protein. For protein folding pattern recognition, PFP-FunD SeqE tool [47] was used. Detection of coiled-coil conformation within the protein was performed using COILS server [48].

### Protein-protein interaction analysis

Protein residues interact with each other for their accurate functions. STRING database [49], known for the prediction of protein-protein interactions, search was performed to identify the possible functional interaction networks of the selected hypothetical protein of *L. acidophilus* bacteria (TDB29877.1).

### Secondary structure prediction

The retrieved protein sequence was used for the prediction of the secondary structure of the hypothetical protein. SOMPA server [50, 51] was used in this regard. The secondary structure prediction was also further cross-checked and validated by SABLE [52] and PSIPRED servers [53].

### Three-dimensional (3D) prediction

The three-dimensional (3D) structure prediction of the hypothetical protein was performed using HHpred server [54, 55] of the Max Planck Institute for Development Biology, Tubigen based on pairwise comparison profile of Hidden Marcov Models (HMMs). Visualization of the 3D structure obtained from the HHpred server was then done by PyMOL program [56]. SWISS-MODEL interactive workspace [57] was also further used to verify the prediction of 3D structure of hypothetical protein by automated comparative modeling. Later, the 3D structure was further refined through YASARA energy minimization server [58] and YASARA view software.

### Quality assessment of the model

Several assessment tools were used for the quality assessment of the predicted 3D structure of the hypothetical protein. PROCHECK, VERIFY3D and ERRAT tools of SAVES server [59–61] were used for the quality assessment of the build model.

### Active site detection

Determination of active site of the hypothetical protein was done using Computer Atlas of Surface Topography of Protein (CASTp) server [62, 63] which provides an online resource for locating, delineating and measuring concave surface regions on three-dimensional structures of proteins.

## Results

### Retrieval of hypothetical protein sequence

The selected a hypothetical protein (TDB29877.1) from *L. acidophilus* is a gram positive bacteria. It contains 606 amino acid residues. Additional information collected from the NCBI database regarding this hypothetical protein is given in Table 1.

**Table 1.** Primary information of the selected hypothetical protein

| Protein individualities | Hypothetical protein information |
| --- | --- |
| Locus | TDB29877 |
| Definition | hypothetical protein E1P27_08605 [*Lactobacillus acidophilus*] |
| Accession number | TDB29877 |
| Version | TDB29877.1 |
| Organism | *Lactobacillus acidophilus* |
| Source strain | PNW3 |
| Host | *Sus scrofa domesticus* |
| Country and collection time | South Africa: Pretoria; Jun-2012 |

### Physicochemical properties analysis

The PortParam tool of the ExPaSy server was used to retrieve the physicochemical properties of the uncharacterized protein. The most abundant amino acid residue observed was lysine (9.7%), followed by leusine (9.6%), aspartic acid (8.6%), glutamine (7.8%) and isoleucine (7.3%). The lowest number of amino acids were cysteine (0.3%), tryptophan (1.2) and histidine (1.5%). Other physicochemical properties of the protein are given in Table 2.

**Table 2.** Physicochemical properties of the hypothetical protein

| Properties | Value |
| --- | --- |
| Molecular Formula | C<sub>3274</sub>H<sub>5024</sub>N<sub>838</sub>O<sub>961</sub>S<sub>12</sub> |
| Molecular weight | 71885.66 |
| Theoretical pI | 5.59 |
| Total number of negatively charged residues (Asp+Glu) | 99 |
| Total number of positively charged residues (Arg+Lys) | 88 |
| Instability index | 38.21 |
| Aliphatic index | 84.80 |
| Grand average of hydropathicity (GRAVY) | -0.636 |

### Homology identification

BLASTp program was used against non-redundant protein sequences and UniProt databases to find out the homologous proteins with having structural similarities to the selected hypothetical protein. The result of BLASTp program were given in Tables 3 and 4.

**Table 3.** Similar proteins obtained from non-redundant protein sequences (nr) database

| Accession No | Organism | Protein name | Score | Ident. % | E-value |
| --- | --- | --- | --- | --- | --- |
| WP_003546145.1 | *Lactobacillus acidophilus* | TerB N-terminal domain-containing protein | 1238 | 100 | 0 |
| WP_170089063.1 | *Lactobacillus amylovorus* | TerB N-terminal domain-containing protein | 801 | 64.76 | 0 |
| WP_052542817.1 | *Lactobacillus sp. OTU4228* | TerB N-terminal domain-containing protein | 796 | 64.27 | 0 |
| WP_202017233.1 | *Lactobacillus kitasatonis* | TerB N-terminal domain-containing protein | 790 | 64.05 | 0 |
| WP_007125014.1 | *Lactobacillus ultunensis* | TerB N-terminal domain- containing protein | 774 | 63.39 | 0 |
| WP_098044344.1 | *unclassified Lactobacillus* | MULTISPECIES: TerB N-terminal domain-containing protein | 759 | 61.83 | 0 |
| WP_060462377.1 | *Lactobacillus crispatus* | TerB N-terminal domain-containing protein | 759 | 59.87 | 0 |
| WP_204781356.1 | *Lactobacillus gallinarum* | TerB N-terminal domain-containing protein | 757 | 61.79 | 0 |

**Table 4.** Similar proteins obtained from UniProt database

| Accession No | Organism | Protein Name | Score | Ident. % | E-value |
| --- | --- | --- | --- | --- | --- |
| A0A4V3BIJ5_9LACO | *Lactobacillus crispatus* | TerB_N domain-containing protein | 1,163 | 62.40 | 1.00E-152 |
| A0A0U5K922_LACDE | *Lactobacillus delbrueckii*<br>*subsp. Bulgaricus* | TerB-N/TerB-C domain | 739 | 32.30 | 1.60E-85 |
| A0A0R2D8P4_9LACO | *Lactobacillus taiwanensis* DSM 21401 | TerB_N domain-containing protein | 663 | 34.20 | 1.30E-76 |
| A0A1G6BEX5_9FIRM | *Ruminococcaceae*<br>*bacterium* FB2012 | TerB-C domain-containing protein | 629 | 29.60 | 8.70E-70 |
| A0A1H9EUD7_9FIRM | *Butyrivibrio* sp. TB | Predicted DNA-binding protein, MmcQ/YjBR family | 540 | 27.00 | 3.70E-56 |
| A0A315Y7B5_RUMFL | *Ruminococcus flavefaciens* | TerB-like protein | 523 | 29.40 | 1.60E-55 |
| A0A3D5MZF8_9FIRM | *Erysipelotrichaceae*<br>*bacterium* | TerB_N domain-containing protein (Fragment) | 492 | 27.10 | 8.20E-51 |
| A0A562SBM9_9SPIO | *Treponema putidum* | TerB-like protein | 475 | 27.60 | 2.30E-48 |
| A0A2M8Z4U1_9FIRM | *Clostridium* *celerecrescens 18A* | TerB-like protein | 433 | 28.90 | 2.10E-42 |

### Multiple sequence alignment and phylogeny analysis

Sequences obtained from BLASTp program and the query sequence (TDB29877.1) were aligned by MUSCLE using MEGA software is shown in Fig 1. Multiple sequence alignment was also cross-checked by Clustal Omega program of EMBL-EBI (S1 File). For the confirmation of homology assessment between the proteins, down to the complex and subunit level, phylogenetic analysis was also carried out using Phylogeny.fr server. One click method was applied to construct the phylogenetic tree on the basis of BLASTp result and multiple sequence alignment which given the similar concept about the query protein (Fig 2).

**Fig 1.**
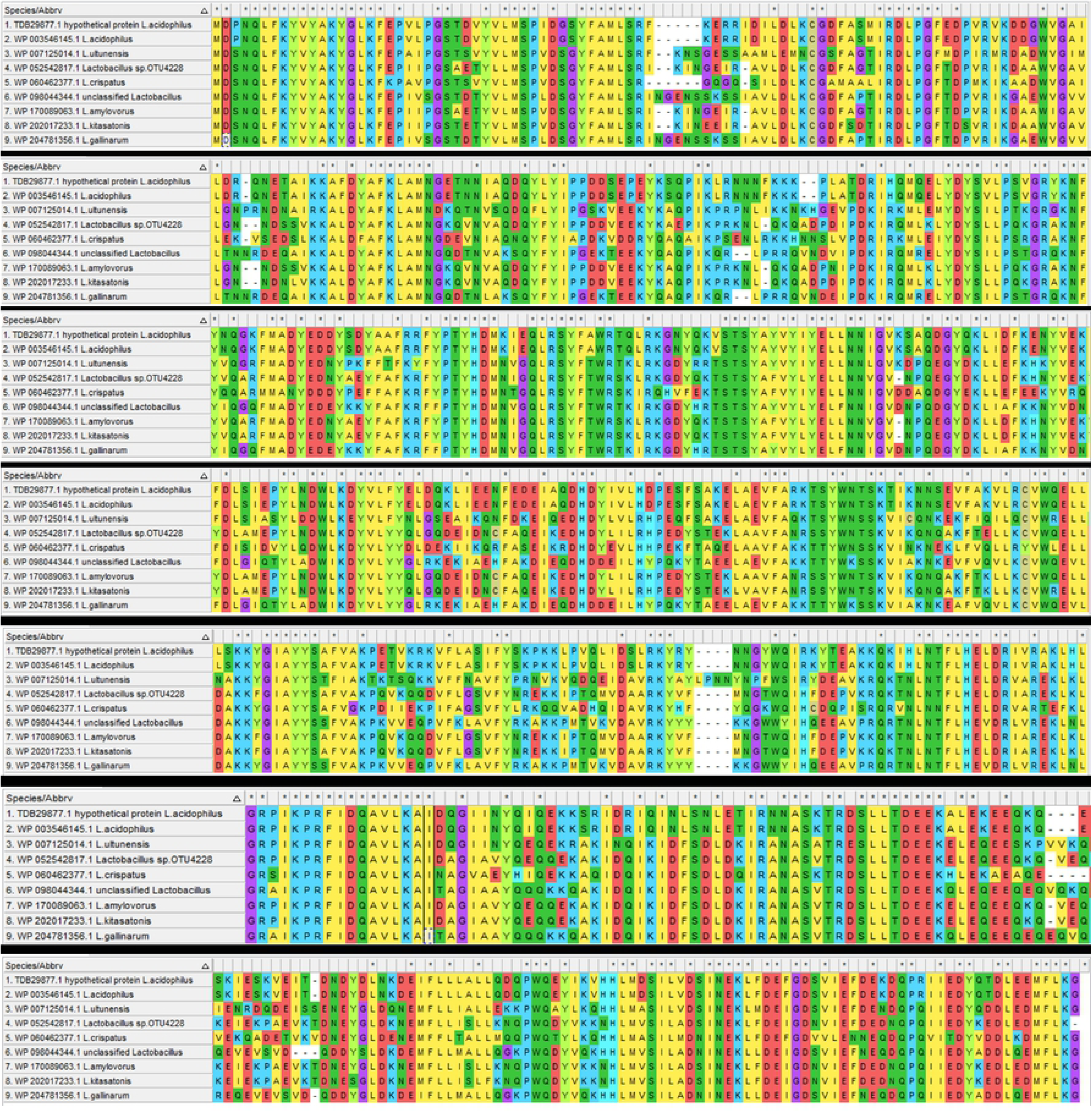
Multiple sequence alignment of different homologous proteins aligned by MUSCLE

**Fig 2.**
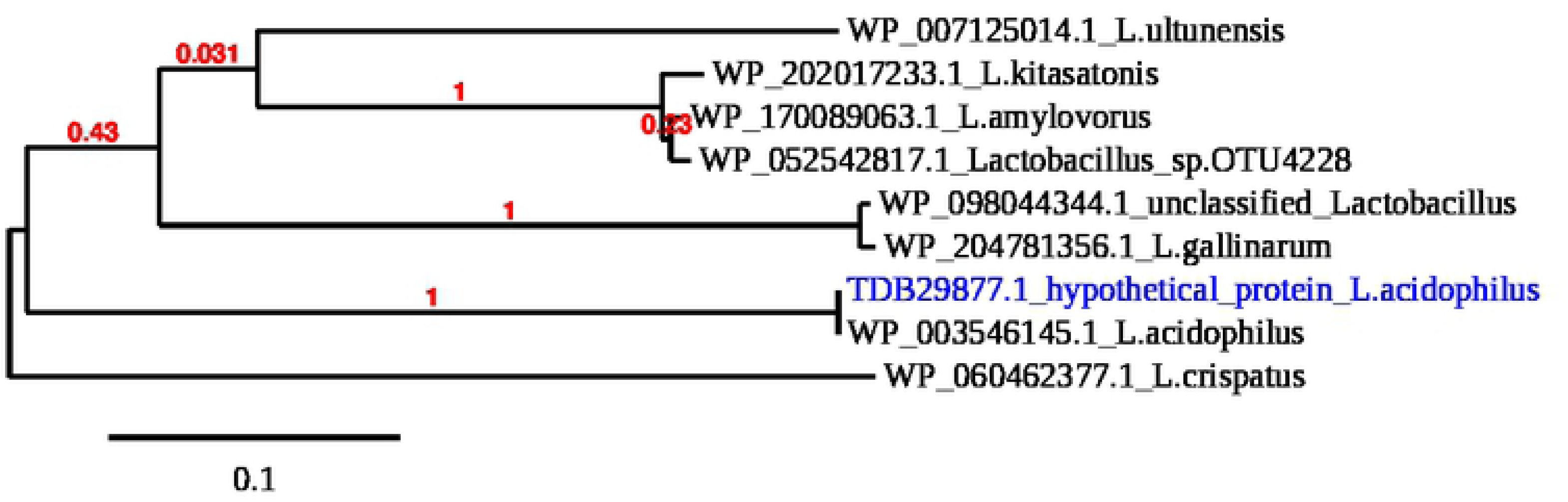
Phylogenic tree with bootstrap confidence values of different proteins from *Lactobacillus* organism

### Subcellular localization analysis

CELLO server was used to identify the subcellular localization of the selected uncharacterized protein. It was found that it’s a cytoplasmic protein. The result obtained from the other servers (PSORTb, SOSUI, TMHMM, HMMTOP and CCTOP) were also revealed the similar indication (Table 5).

**Table 5.** Subcellular localization of the hypothetical protein

| Subcellular localization analysis | Result |
| --- | --- |
| CELLO 3.0 | Cytoplasmic |
| PSORTb | Cytoplasmic membrane |
| SOSUI | Soluble protein |
| TMHMM 2.0 | No transmembrane helices present |
| HMMTOP | No transmembrane helices present |
| CCTOP | Not transmembrane protein |

### Functional annotation analysis

The conserved domain search (CD-search) revealed that (shown in Fig 3) the selected hypothetical protein had two domains, TerB-N terminal domain (accession no: pfam 13208) and TerB-C domain (accession no: cl21414). The TerB-N domain is found N-terminal to TerB, and TerB-C containing proteins. TerB-C occurs in the C terminus of TerB in TerB-N containing proteins. Pfam server predicted the TerB N-terminal domain at 141-360 amino acid residues with an e-value 8.8e-47 and TerB-C domain at 467-599 amino acid residues with an e-value 3.2e-21. Motif and InterProScan servers also forecasted the same domains with at similar alignment position. However, ScanProsite tool of ExPasY server did not find any hit while searching for motif. Superfamily server revealed presence of YjbR-like superfamily. PFP-FunD SeqE tool predicted the fold type of the selected hypothetical protein as Belta-grasp. The x-axis of output graph from COILS server represented the position of the amino acid number in the protein (starting at the N-terminus) and the y-axis showed the coiled coil whereas ‘window’ refers to the width of the amino acid window that is scanned at one time.

**Fig 3.**
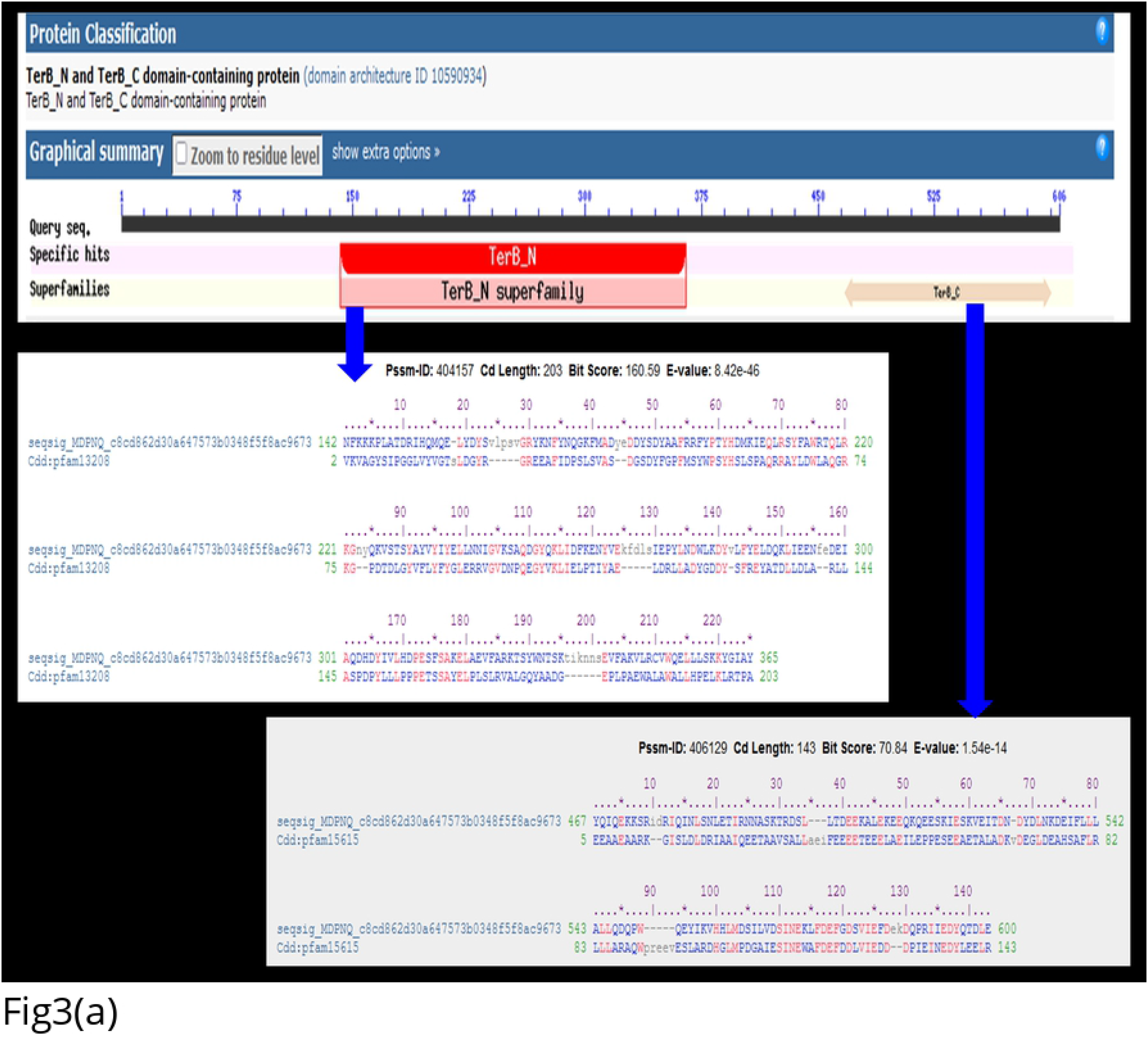
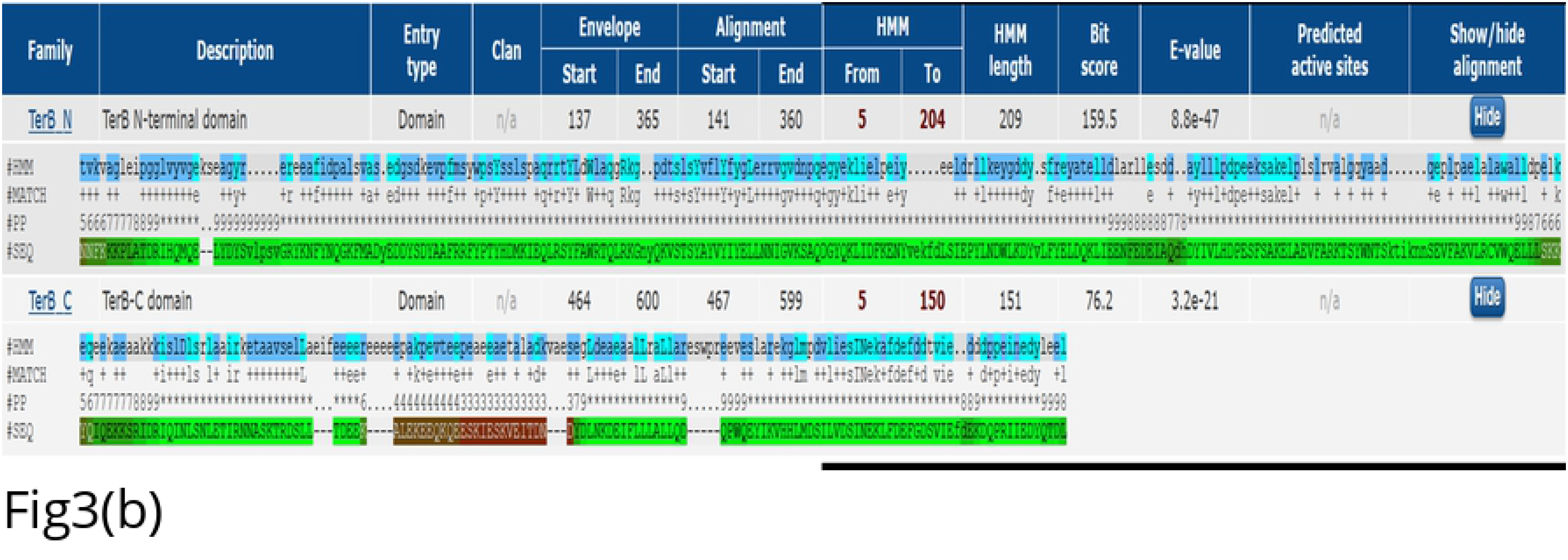
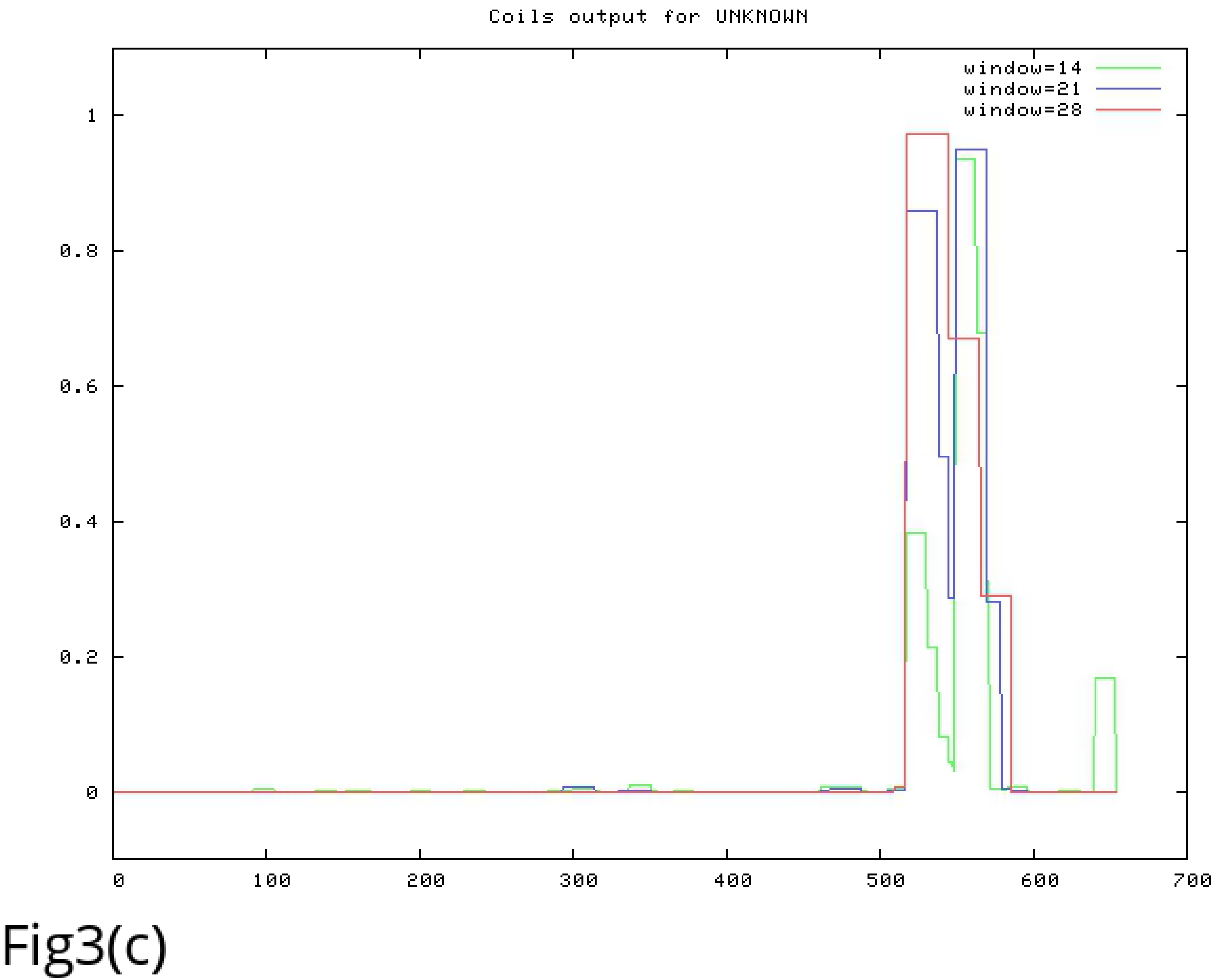
Functional annotation of the hypothetical protein: Fig 3(a) NCBI CD-search result; Fig 3(b) Search result in Pfam server; Fig 3(c) Result of COILS server: coil shows the heptads corresponding to the residue window 14 (green), 21 (blue) and 28 (red)

### Protein-protein interaction analysis

STRING database was used to analyze the protein-protein interaction. The result obtained from the STRING server revealed that the query protein interacted with other functionally known and unknown or uncharacterized proteins (Fig 4). The selected hypothetical protein of *L. acidophilus* organism (TDB29877.1) showed a high confidence interaction with LBA0469 and LBA0470 protein (same score 0.979) followed by LBA0471 (score 0.845), LBA0466 (score 0.550), LBA0110 (score 0.478), amtB (score 0.464), PspC (score 0.458) and LBA1740 (score 0.418). Of them, there are one COG1201 Lhr-like helicases, one ammonium transport protein, one surface protein PspC and one putative membrane protein. One protein is from Cytochrome P450 71C1 and annotation of three proteins are not available yet.

**Fig 4.**
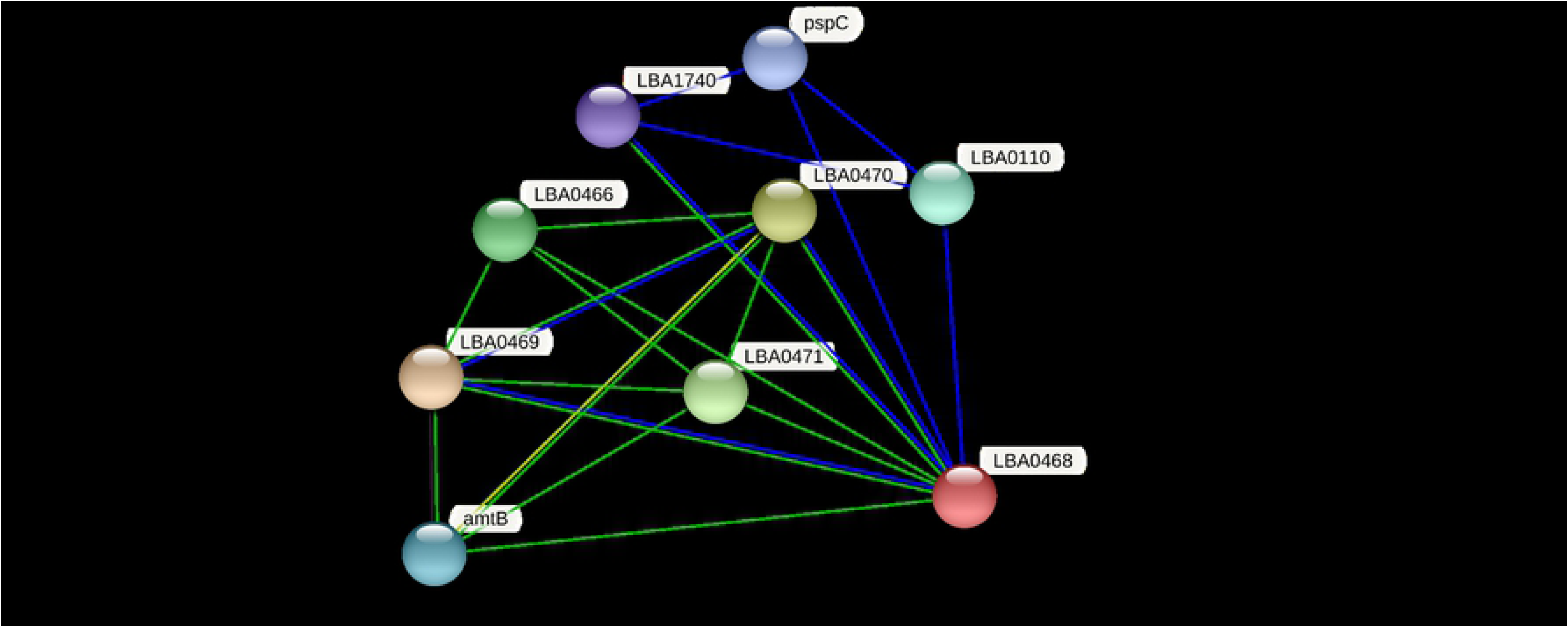
STRING network analysis of the hypothetical protein, indicates as LBA0468

### Secondary structure prediction

Prediction about the secondary structure of the hypothetical protein which includes α-helices, β-sheets, extended strands, turn and coils were obtained from the SOMPA, SABLE and PSIPRED servers (Fig 5). The result of predicted secondary structure of the hypothetical protein from SOMPA server showed that alpha-helices were most predominant (50%) followed by random coil (36.8%), extended strand (10.73%) and beta-turn (2.48%). Similar type of outputs were obtained while validating the secondary structure using SABLE and PSIPRED tools.

**Fig 5.**
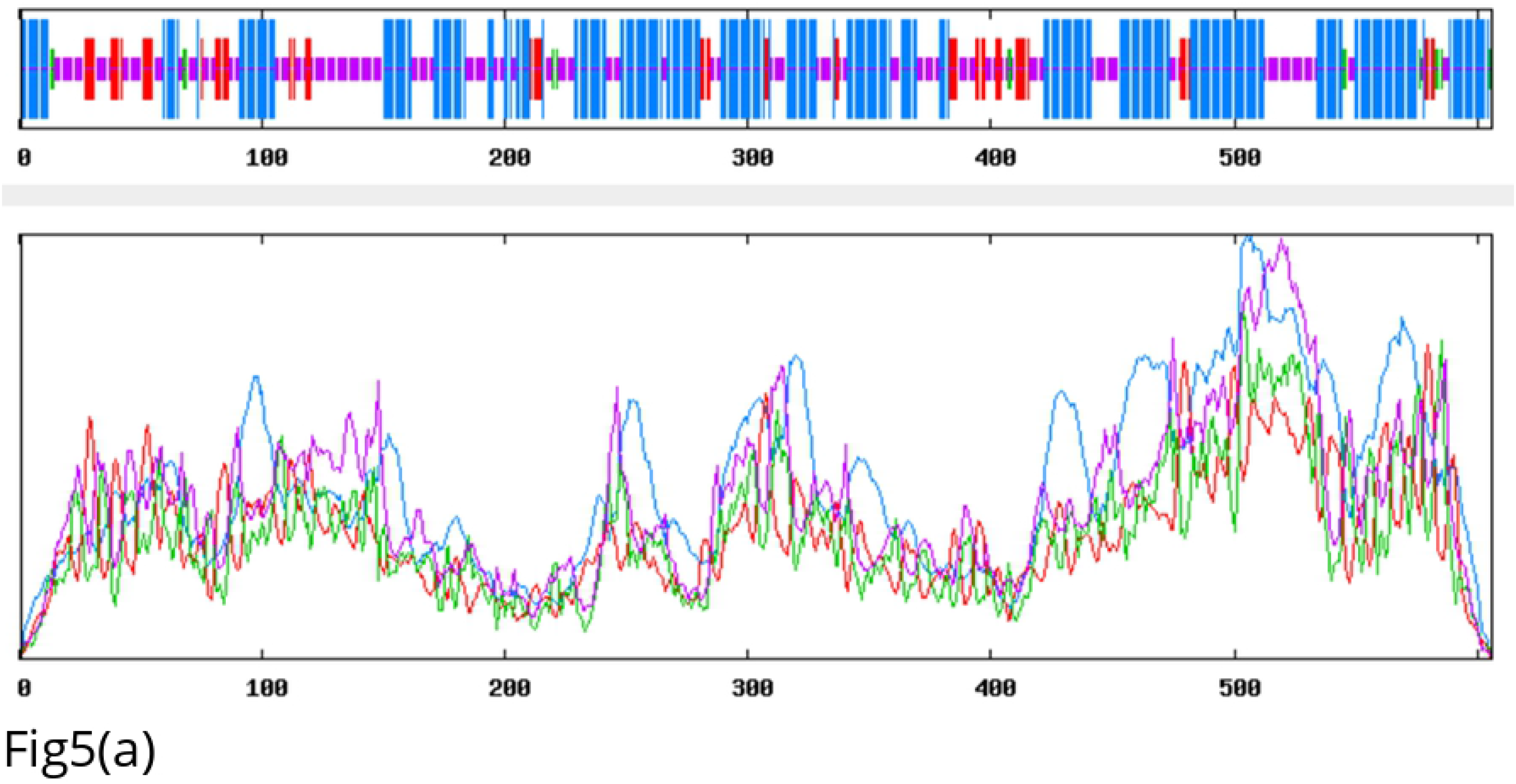
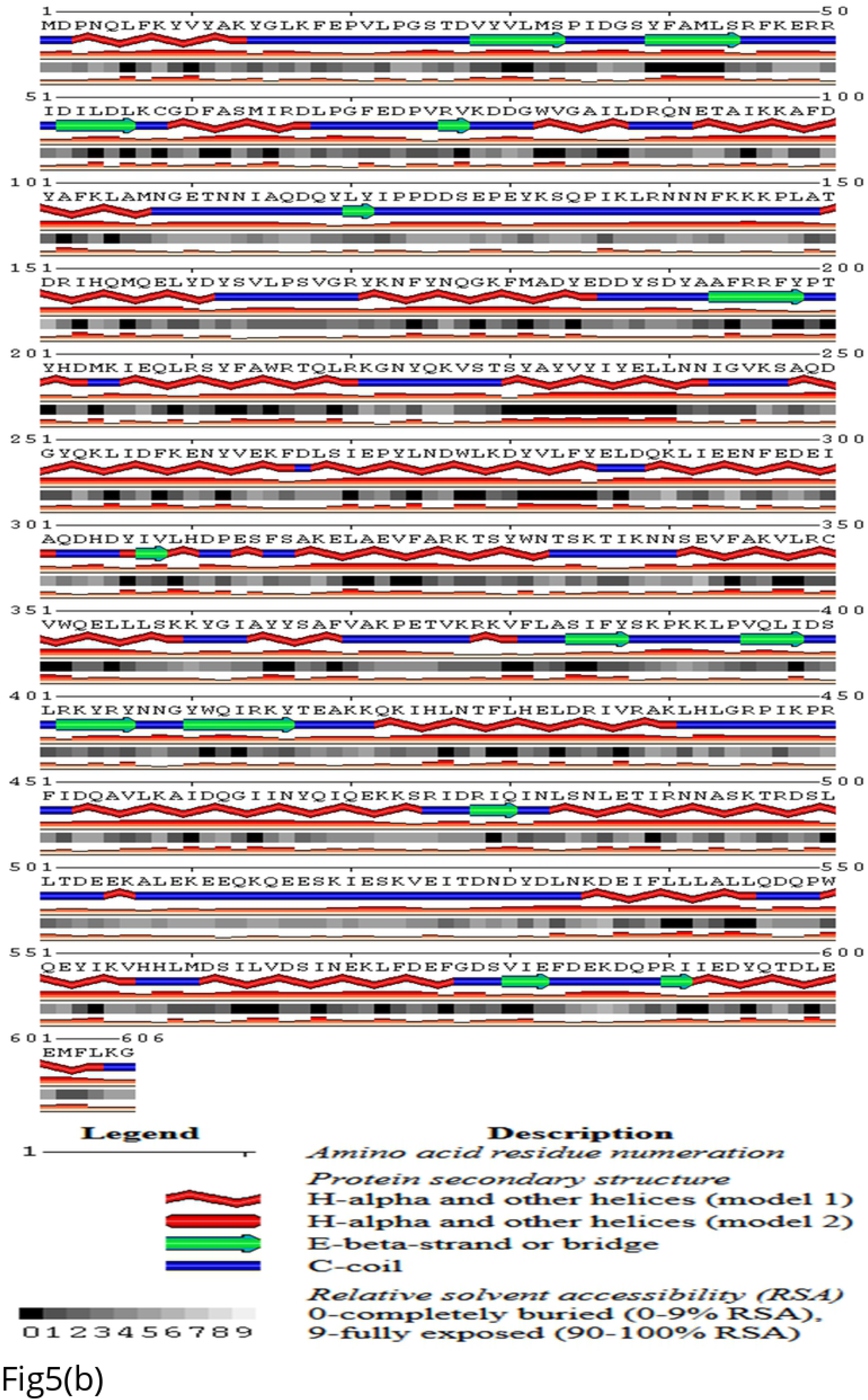
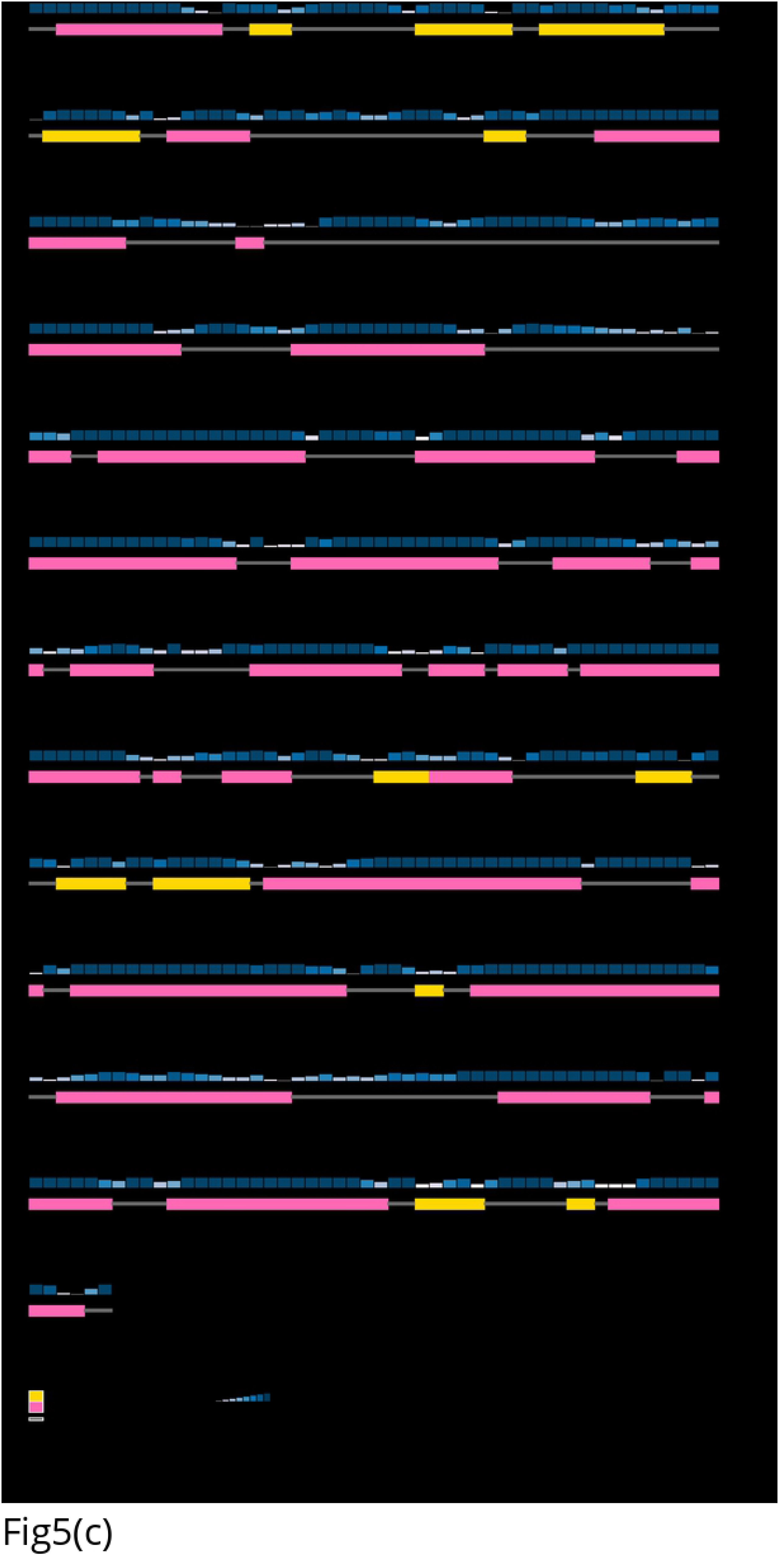
Secondary structure of hypothetical protein predicted by-Fig 5(a) SOMPA server (The window width, similarity threshold and number of states were 17, 8 and 4 respectively); Fig 5(b) SABLE server; Fig 5(c) PSIPRED server

### Three-dimensional (3D) prediction

Prediction of the three-dimensional (3D) structure of hypothetical protein was done by using HHpred server. This server predicted 3D structure of the protein (Fig 6) having 99.24% identity with the highest scoring template (PDB ID: 3H9X_A). 3H9X_A is a crystal structure of the PSPTO_3016 protein from *Pseudomonas syringae* organism with four chains (Chain A, B, C and D). Further validation of the 3D structure prediction by SWISS-MODEL interactive workspace revealed that the oligo-state of the protein is a monomer [64]. The crystallographic resolution of the template used to the model protein was 2.51Å by adopting the X-ray diffraction method [65]. Global quality estimate, local quality estimate, comparison of protein size residue and model template alignment were also explored from this server (Fig 7). Later, the 3D structure was further modified by YASARA energy minimization server. The energy calculated before energy minimization was −10988.9 kJ/mol whereas it was reduced to −55991.2 kJ/mol after energy minimization. The initial score was −2.99 while the final score was −0.19 after energy minimization.

**Fig 6.**
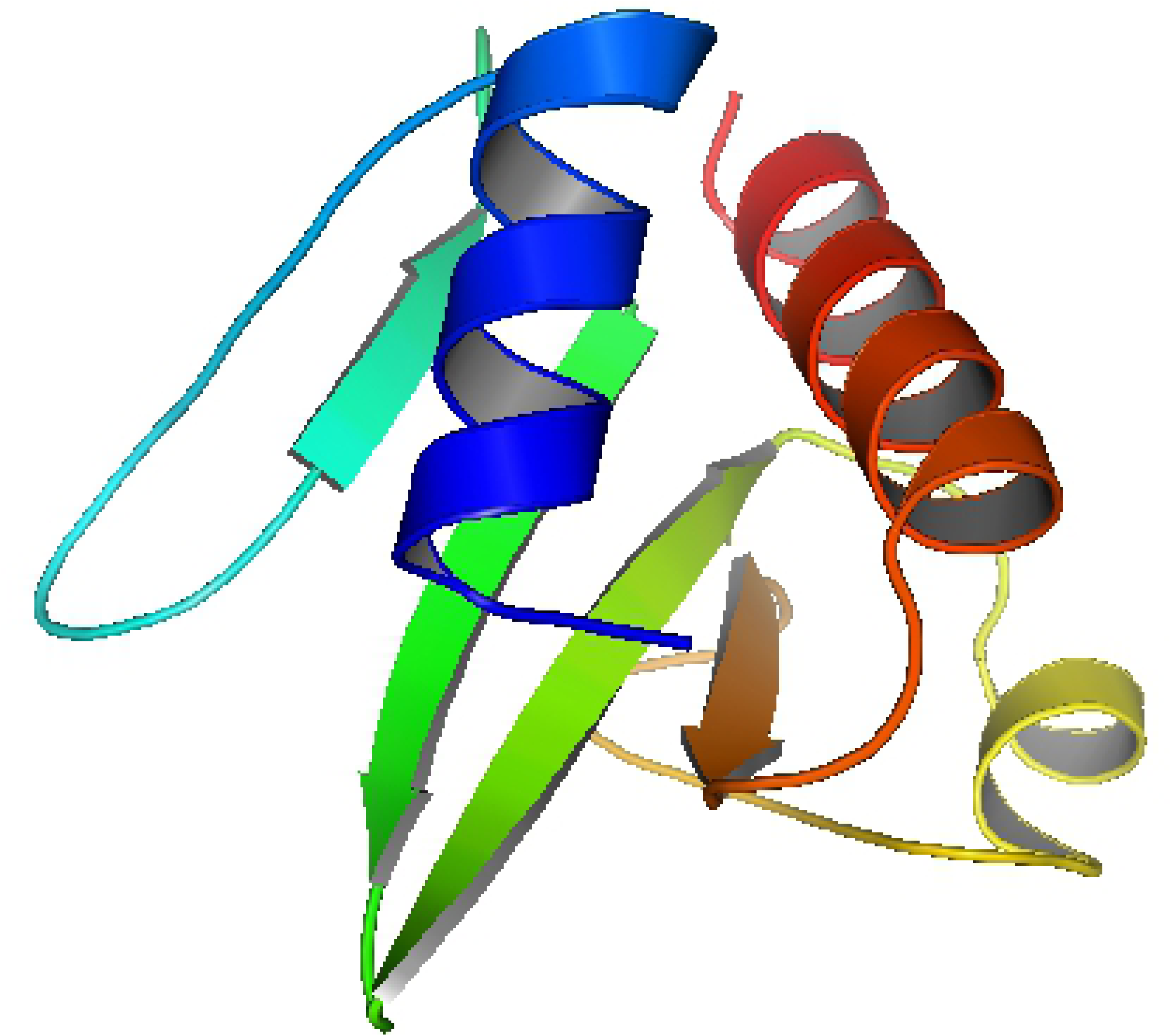
Predicted 3D structure of the hypothetical protein

**Fig 7.**
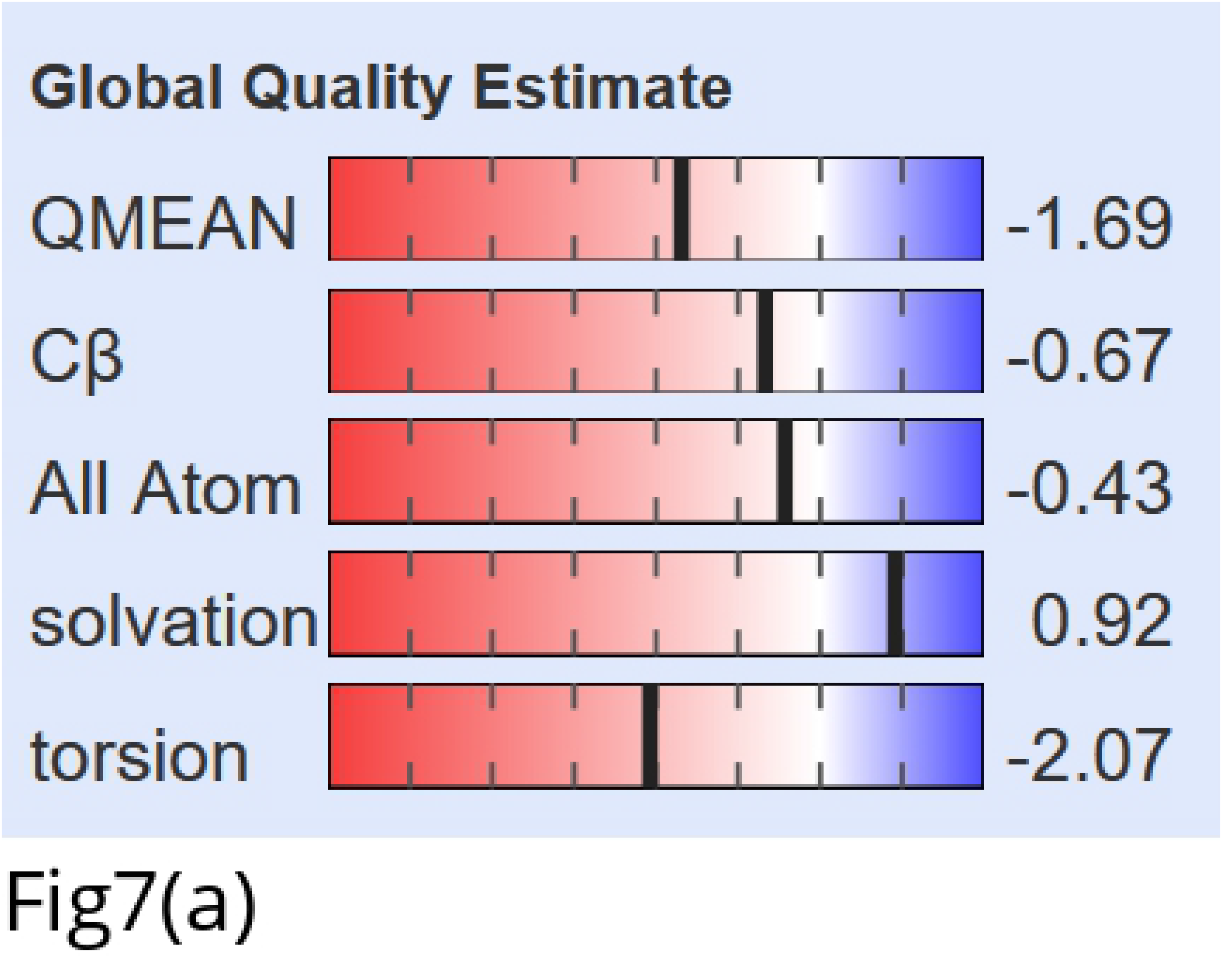
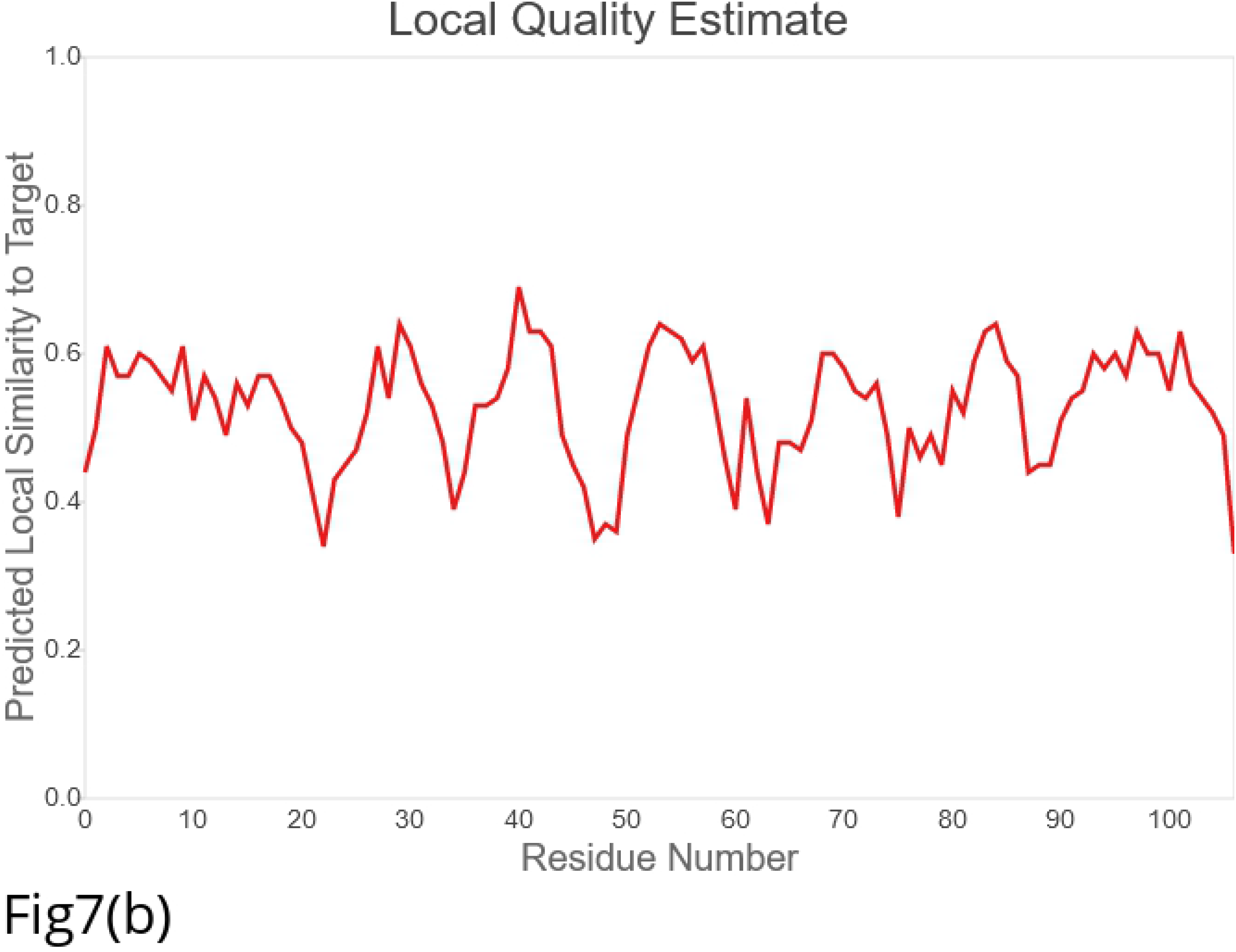
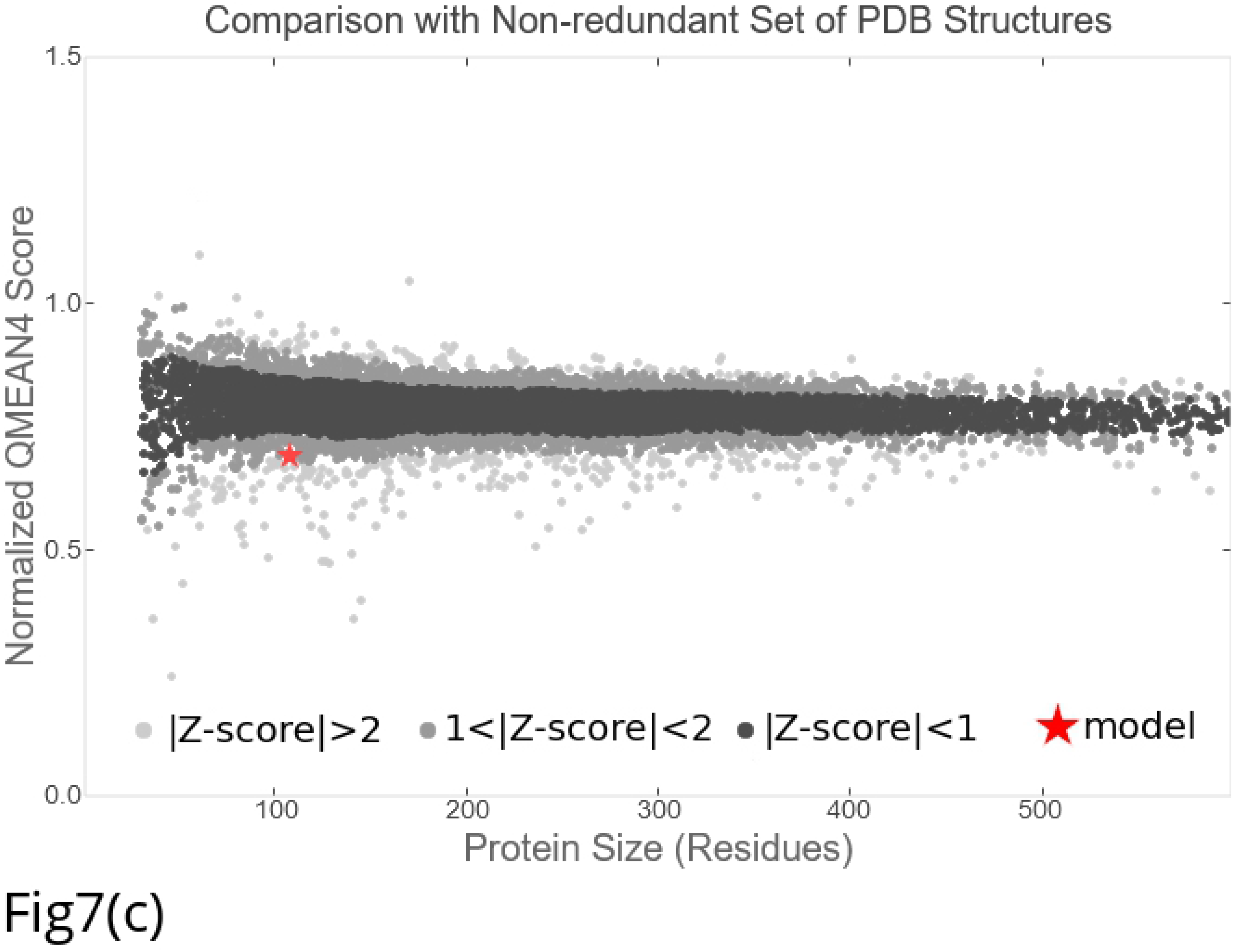
The assessment of 3D structure using SWISS-MODEL interactive workspace: Fig 7(a) Global quality estimate; 7(b) local quality estimate; Fig 7(c) comparison of the protein size residue

### Quality assessment of the model

Validation of 3D structure of the hypothetical protein was done through several quality assessment steps. Assessment of the 3D model was done by PROCHECK tool through Ramachandran plot analysis (Fig 8), where the distribution of φ and ψ angle in the model within the limits were shown. This result also showed that residues in the most favored regions covered 92.4% (Table 6). Then, the structure again verified by VERIFY 3D and ERRAT tools and found 90.65% of the residues had average 3D-1D score ≥ 0.2 and overall quality factor was 72.72 respectively.

**Fig 8.**
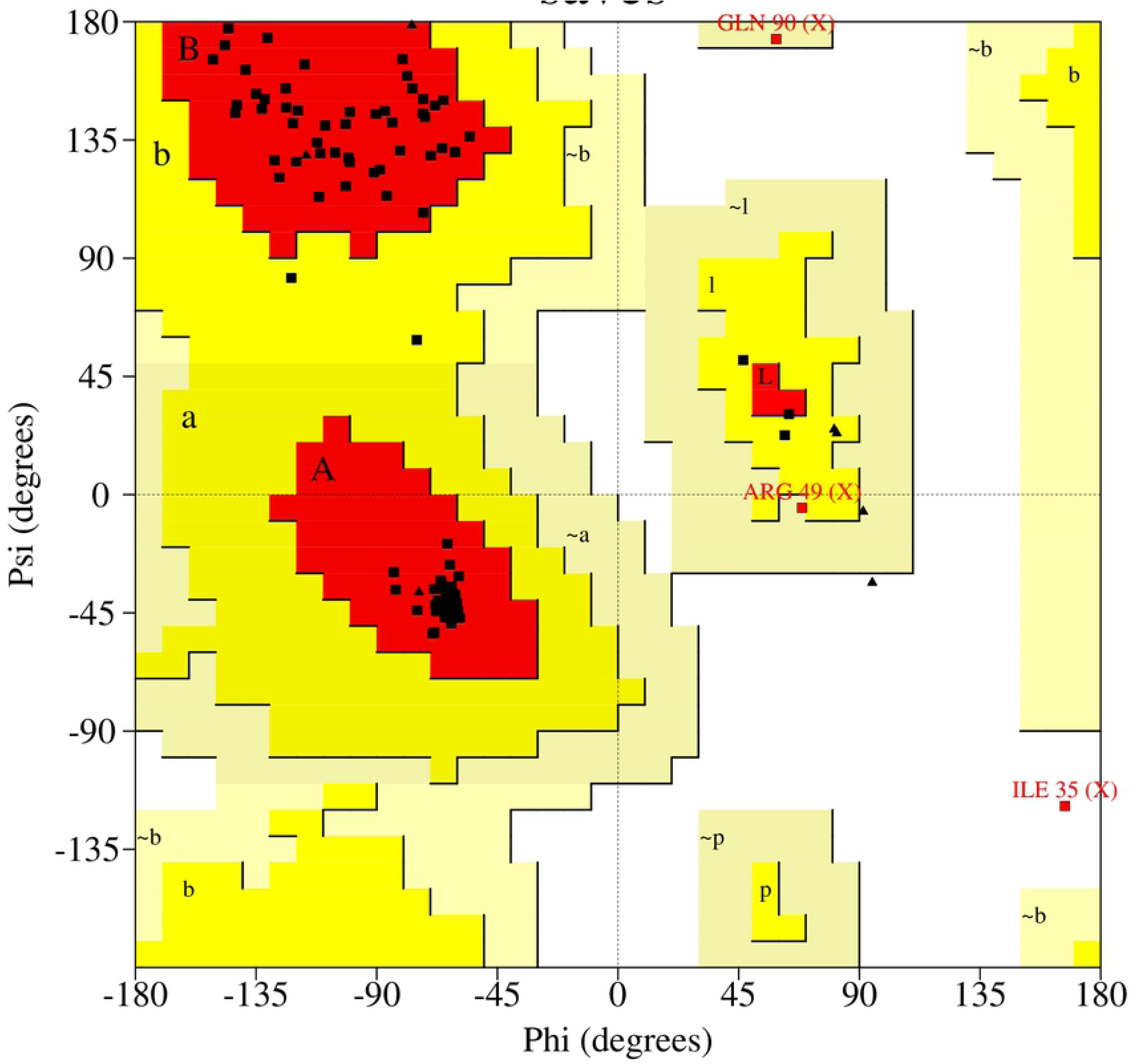
Ramachandran plot for the 3D model of the hypothetical protein validated by PROCHECK program

**Table 6.**
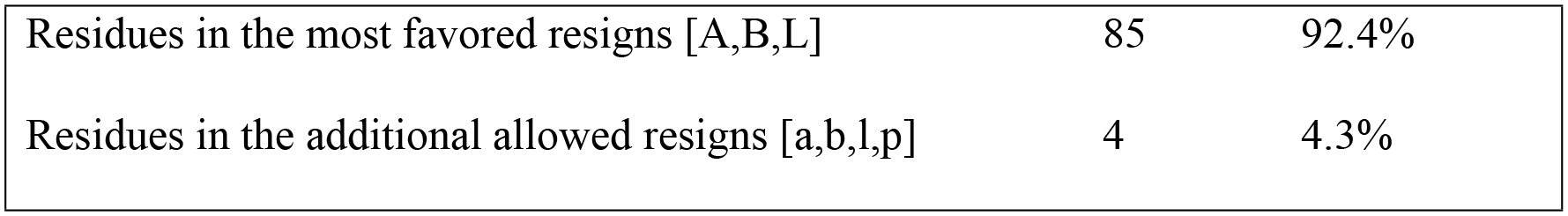
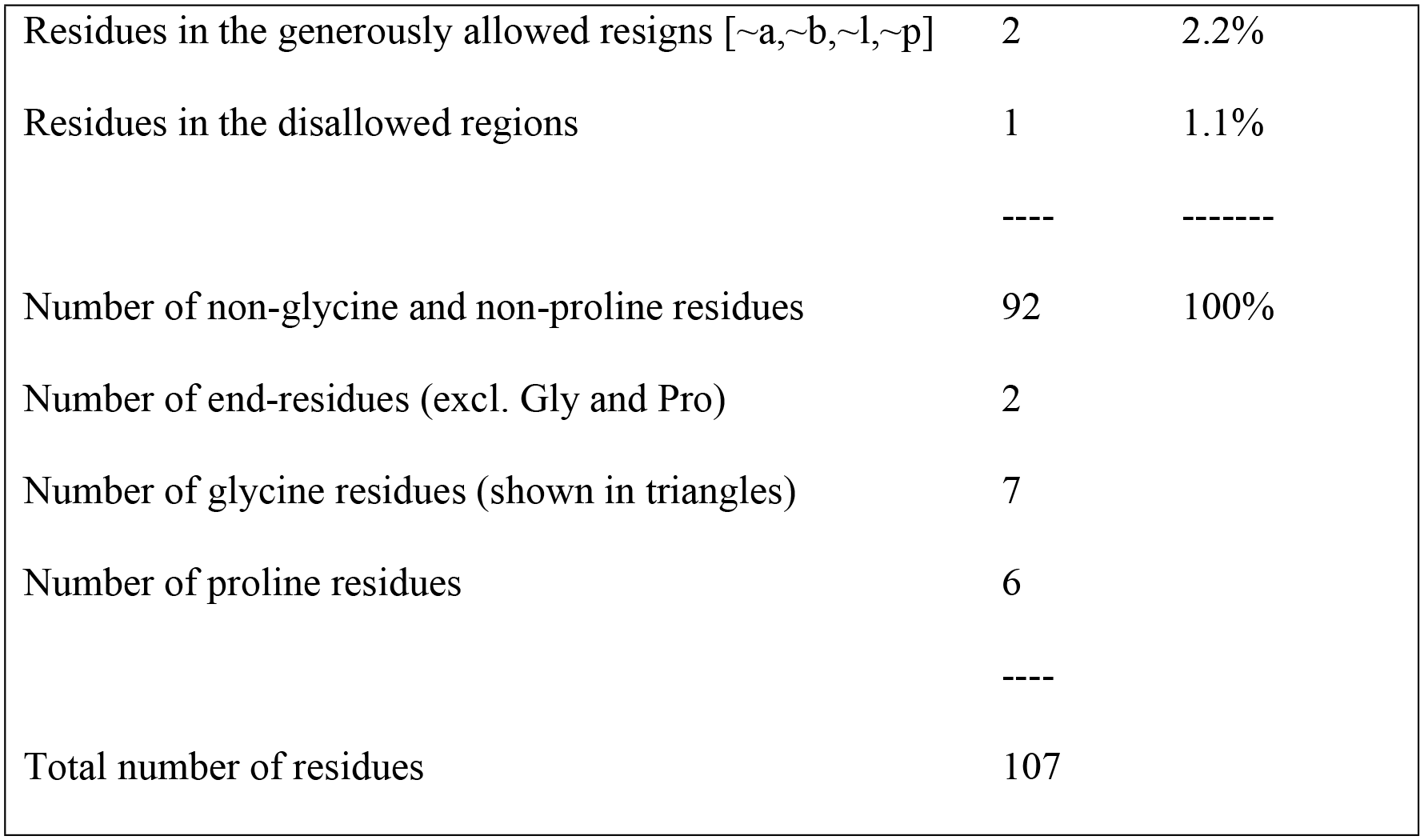
Ramachandran plot statistics of the hypothetical protein

### Active site detection

Computer Atlas of Surface Topography of Protein (CASTp) server was used to determination the active sites with the amino acid residues of the hypothetical protein (Fig 9). The result from CASTp calculation revealed a total of 18 active pockets of the hypothetical protein. The best active site found in the areas (SA) with 126.75 and a volume (SA) of 78.13 amino acids.

**Fig 9.**
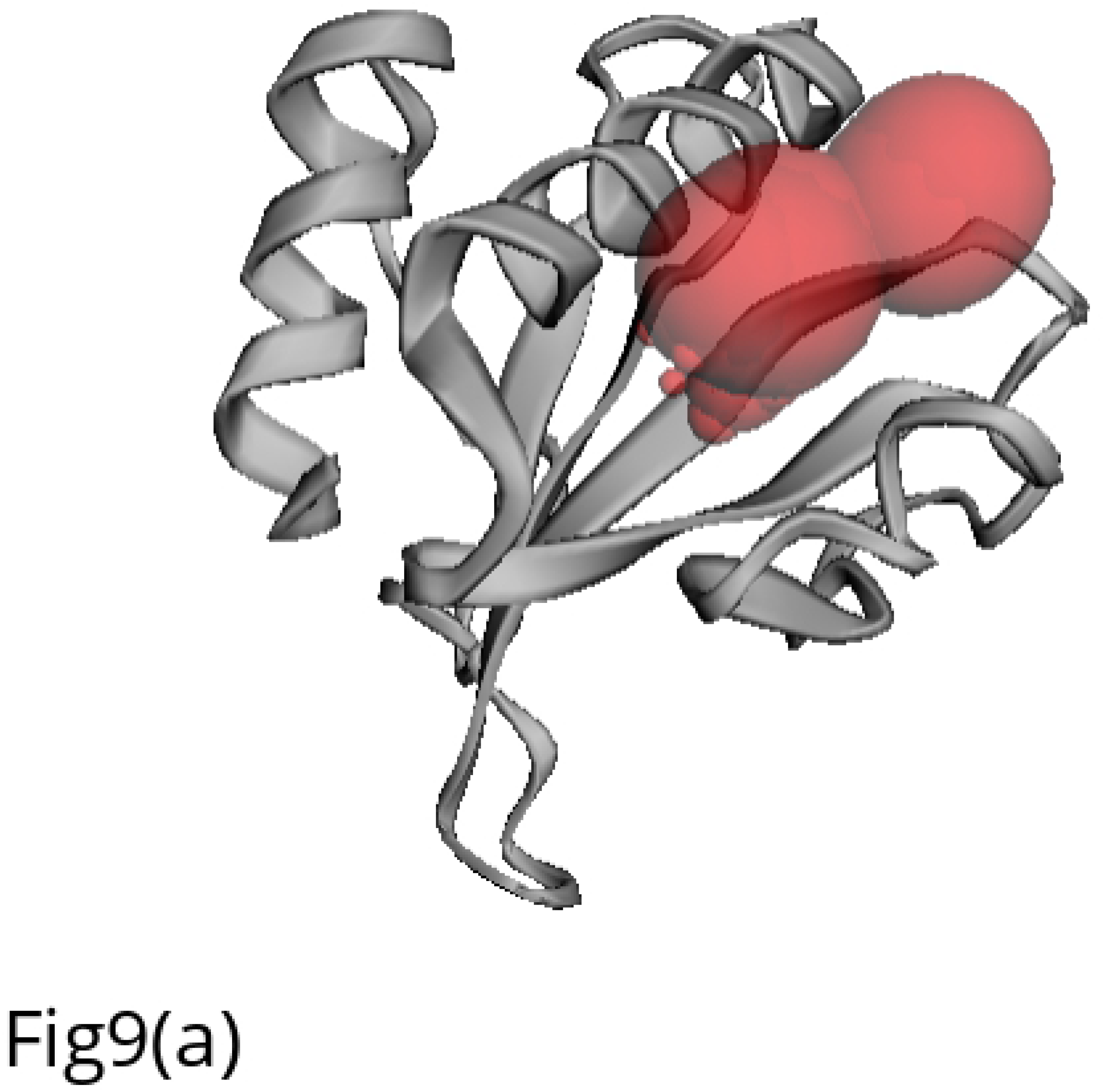
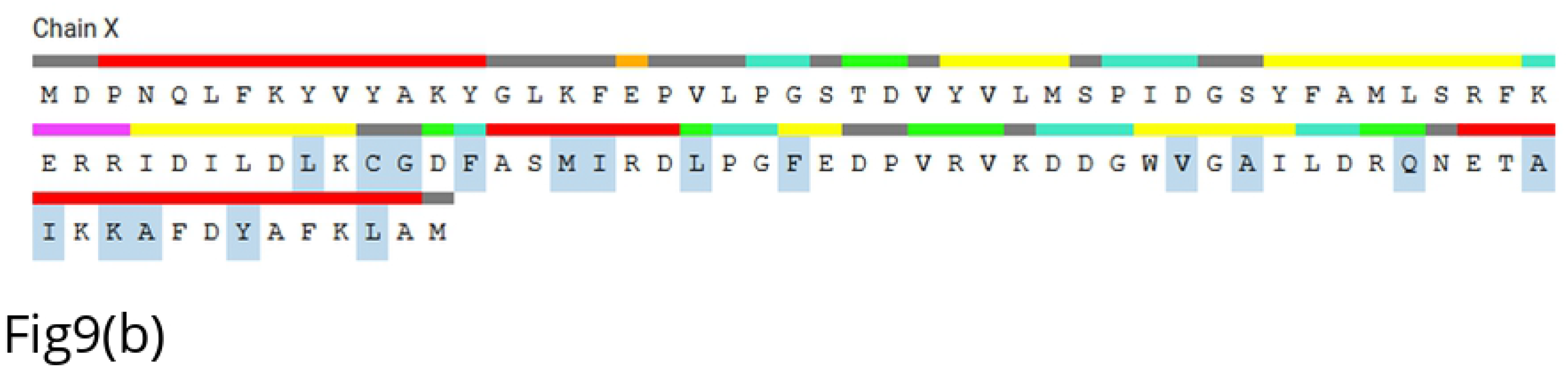
Active site detection of the hypothetical protein using CASTp: Fig 9(a) The red sphere indicates the active site of the protein; Fig 9(b) Sequence of active amino acid residues of the largest pocket

## Discussion

Since the *L. acidophilus* bacteria are well known for its beneficial properties and are being used as probiotics, a hypothetical protein from this organism have targeted to examine it’s involvement in the defensive mechanism against pathogenic organisms. Thus, the amino acid sequence of the targeted hypothetical protein was retrieved for further investigation. The selected hypothetical protein had a total of 606 amino acids. The computed instability index (38.21) classifying the protein as stable one because instability index value below 40 indicates a protein as stable and above 40 indicates as unstable [66]. The selected protein’s aliphatic index (94.34) indicated it’s stability over a wide range of temperature and the negative GRAVY value (−0.636) indicated the hydrophilicity nature of the hypothetical protein [67].

Homology analysis revealed that the query protein has structural similarities with other TerB- N and TerB-C domain containing proteins from various *Lactobacillus* species. Multiple sequence alignment using MUSCLE and Clustal Omega program produced alignments between the selected sequences from BLASTp using seeded guide trees and HMM profile-profile techniques. The phylogenetic tree displayed the highest degree of similarity between the studied hypothetical protein and its related proteins for homology modeling obtained by BLASTp of NCBI. The bootstrapping confidence levels of the analysis stated the closed similarity between the query protein (TDB29877.1) and TerB N-terminal domain-containing protein (WP_003546145.1) of *L. acidophilus* organism. Other homologous proteins from various *Lactobacillus* species formed separate clades having varied structural similarity with the studied hypothetical protein.

Basically, subcellular localization of protein indicates where the protein resides in a cell; it may be in outer membrane, inner membrane, periplasm, extracellular or in cytoplasm [68]. The functional properties, interaction and genome annotation are highly influenced by its subcellular localization. Results from CELLO and PSORTb servers indicated the hypothetical protein as a cytoplasmic one. SOSUI server was also depicted the selected protein as a soluble protein. Absent of transmembrane helices predicted by TMHMM and HMMTOP also emphasized the result of being cytoplasmic protein. In addition, CCTOP server also summarized that the query protein was not a transmembrane protein, thus it’s a cytoplasmic protein. Such subcellular identification analysis indicated that the hypothetical protein might be involved in recovering disease state through discovering some novel drugs [69]. As the membrane protein can be used as a potential vaccine target and the cytoplasmic proteins may act as promising drug targets [70], the selected hypothetical protein may be a good source for producing various beneficial pharmaceutical products or healthy food items. The response against chemical stress and anti-viral defense systems of bacteria are constituted by the Ter gene products. The TerB_N has a predominantly alpha-helical structure and contains an absolutely conserved glutamate. The presence of a conserved acidic residue suggested that it might chelate metal like TerB. These proteins occur in a two-gene operon containing an AAA+ ATPase and SF-II DNA helicase suggesting a role in stress-response or phage defense [71]. TerB-C domain also displays multiple conserved acidic residues. The presence of conserved acidic residues in both TerB-N and TerB-C suggested that they, like the TerB domain, might also chelate metals. These two domains might also occur together in the same protein independently of TerB [71]. Motif, Pfam and InterProScan servers also confirmed the presence of TerB-N and TerB-C domains in the selected hypothetical protein. YjbR-like superfamily of the hypothetical protein is expected to contain the DNA binding domain comprising the ‘double wing’ motif [72]. Moreover, protein fold plays a significant role in their function and hence the fold prediction has also been applied in order to further validate the predicted function. PFP-FunD SeqE tool revealed the fold type of the hypothetical protein as ‘Belta-grasp’ which indicated that the protein might play role in hydrolase activity [73].

The function of a target protein and drug availability of molecules can be predicted by analyzing protein-protein interaction [49]. Protein-protein interaction network analysis showed that the query protein (TDB29877.1, shown as LBA0468 in Fig 4) highly interacted with proteins LBA0469, LBA0470 and LBA0471 within the network; they are also neighborhood in the genome. These interactions give an indication about the selected hypothetical protein that it might be involved in iron ion binding, DNA and RNA metabolisms and numerous repair mechanisms that maintain cellular integrity [29, 74].

It was obtained from SOMPA saver prediction that the selected secondary structure of hypothetical protein was an alpha-helices dominating protein. The window width, similarity threshold and number of states were 17, 8 and 4 respectively. Confidence of prediction from PSIPRED server also stated alpha-helices dominating output. In addition, SABLE server forecasted the secondary structure of the protein having a good confidence of prediction.

The HHpred server forecasted a 3H9X_A protein template with highest score (106.27). 3H9X belongs to the protein Pspto_3016 of *Pseudomonas syringae*. Pspto_3016 is a 117-residue member of the protein domain family PF04237, which is to date a functionally uncharacterized family of proteins [75]. The GMQE (Global Model Quality Estimation) and QMEAN value of the selected model from the SWISS-MODEL interactive workspace analysis were 0.06 and −1.69 respectively [57]. GMQE is a quality estimation which combines properties from the target–template alignment and the template structure and the resulting GMQE score is expressed as a number between 0 and 1. The QMEAN Z-score provides an estimate of the ‘degree of nativeness’ of the structural features observed in the model on a global scale [76, 77] and scores of −4.0 or below are an indication of models with low quality. The GMQE and QMEAN score of the selected model indicated it as a comparatively reliable and better quality model. The comparison plot and model quality scores of individual models are related to scores obtained for experimental structures of similar size. The x-axis shows protein length (number of residues). The y-axis is the normalized QMEAN score. Every dot represents one experimental protein structure. Black dots are experimental structures with a normalized QMEAN score within 1 standard deviation of the mean (|Z-score| between 0 and 1), experimental structures with a |Z-score| between 1 and 2 are grey. Experimental structure that are even further from the mean are light grey [76, 77]. The actual model is represented as a red star meant that our model was within the grey region. In addition, reduced energy and improved score of the predicted model applying YASARA energy minimization tools indicated the model structure as more stable one [58].

Ramachandran plot analysis showed that 92.4% of the residues belonged to the most favored regions. Residues in additional allowed regions and generously allowed regions were 4.3% and 2.2% respectively, which indicated reliability of the model quality. It is generally accepted that more than 90% of the residues in the most favored regions is likely to be a reliable 3D model [78]. The environmental profile or the amino acid environment for non-bonded atomic interactions of the model is good as VERIFY 3D analysis revealed that 90.65% of the residues had average 3D-1D score ≥ 0.2. Overall quality factor obtained through ERRAT was 72.72 which indicated a high quality model. Higher scores indicate higher quality and the generally accepted range is >50 for a high quality model [79].

A probe radius of 1.4Å was used for computing solvent accessible surface area while calculating the active sites of the hypothetical protein using CASTp server. It also measured the exact volumes and areas, as well as sizes of the mouth openings of the active pockets. These metrics were calculated analytically, using both the solvent accessible surface model called Lee and Richards’ surface model [80] and the molecular surface model called Connolly’s surface model [81].

## Conclusion

The current study was designed to forecast the structures and biological functions of a hypothetical protein (TDB29877.1) from *L. acidophilus* bacteria through an *in-silico* approach. All the above findings applying various bioinformatics tools suggested that the selected hypothetical protein from probiotic type bacteria plays role in responding during stress condition or phage defense mechanism. It was also found that the hypothetical protein of interest is cytoplasmic in nature containing ‘Belta-grasp’ fold. These findings may encourage researchers who are interested to work with such beneficial probiotic bacteria to produce various feed additives or healthy food products. Therefore, the outcome of this study in determining structures and functions of the uncharacterized protein indicate reliability of computational approach using bioinformatics tools, thereby assisting experimental validation research on a protein.

## Acknowledgement

The author is grateful to Mr Mohammad Uzzal Hossain of Bioinformatics Division at National Institute of Biotechnology, Bangladesh for providing extraordinary mental support, guidelines and courage to work on such topic.

## Supporting information

**S1 File. Multiple sequence alignment of different homologous proteins aligned by Clustal Omega program of EMBL-EBI**

